# Soil viruses reduce greenhouse gas emissions and promote microbial necromass accrual

**DOI:** 10.1101/2024.03.13.584929

**Authors:** Xiaolong Liang, Shiyu Sun, Yujun Zhong, Ying Zhang, Shuo Wang, Yongfeng Wang, Ninghui Xie, Lu Yang, Mark Radosevich

## Abstract

Viral-induced microbial mortality has been proposed as a major contributor in shaping microbial community structure and function, soil carbon (C) accrual and mobilization of plant available nutrients. Yet, how soil viruses influence soil organic C (SOC) turnover and sequestration remains unknown. Here, we performed microcosm experiments with two distinct soils from grassland (GL) and agricultural (AG) sites and interrogated the roles of soil viruses in driving microbial community succession, SOC transformation and sequestration. The results show that soil viruses affected microbial C use efficiency and reduced respiration in microbial communities obtained from both GL and AG soils. Soil viruses affected microbial successional trajectories (via predation of dominant populations) and functional gene profiles triggering a significant decrease in CO_2_ and N_2_O emissions. The impact of soil viruses on microbial community composition in GL microcosms was much less pronounced compared with that in AG microcosms, suggesting contrasting virus-host interaction patterns under different environmental settings. Viral infection significantly enhanced microbial necromass accumulation thereby increasing SOC and total nitrogen (TN) content. The results implicate viral-mediated microbial mortality as a key factor influencing the distribution of C between mineralization and soil C storage pathways. We proposed ^“^viral loop^”^ to explain the crucial function of soil viruses in SOC turnover and sequestration.

## 1. Introduction

The carbon (C) cycle determines the steady-state level of gaseous forms of C (e.g., CO_2_ and CH_4_) in the atmosphere and is fundamental to global climate. Terrestrial ecosystems represent important C sinks, and large quantities of stable organic C accumulate and are stored in underground soil systems (Bossio et al., 2020). Microorganisms are key drivers of soil C cycling. Microbial growth and activities mediate organic C mineralization releasing CO_2_ into atmosphere, while microbial assimilation of organic compounds and biomass turnover prompt C stabilization via biological pump (Liang et al., 2017; Jansson and Hofmockel, 2020). However, viral predators of soil microorganisms and their impact on transformation and storage of C have just begun to be revealed.

Viruses are the tiniest and most abundant predators in soil and prey on microbes by infecting and ultimately lysing host cells. Viral infection may significantly alter host cell physiology, metabolism, and evolution, which have important implications for soil ecosystem functioning (Liang et al., 2020; Chevallereau et al., 2022; Jansson and Wu, 2022). For example, viral-induced mortality has been proposed as a major driver of microbial community structure, food web dynamics, and biogeochemical cycling (Albright et al. 2022; Wang et al., 2022; Nicolas et al., 2023). Though viruses are known to be the most abundant and active members of soil microbial communities, relatively little is understood concerning their ecological function, especially in biogeochemical processes and soil C accrual.

Studies in marine ecosystems suggest that approximately 10-40% of bacterial biomass is lysed by viruses each day releasing the immense amount of dissolved organic matter (DOM) that becomes readily available for other microorganisms (Fuhrman, 1999; Suttle, 2007; Lara et al., 2017). This is often referred to as the ^“^viral shunt^”^ of the microbial loop and was first proposed by Wilhelm and Suttle (1999). The released nutrients within the microbial loop can stimulate microbial growth, accelerating emissions of CO_2_ into the atmosphere. Besides the DOM that can be rapidly recycled, cell lysates also include sticky cell debris which could form larger polymeric particles leading to aggregation (Weinbauer, 2004). A complimentary notion was then proposed by Weinbauer (2004) suggesting the aggregation creating organic particles exceeding colloidal size sink becoming an export of C to deeper depths. The process that may promote C sequestration was later termed ^“^viral shuttle^”^ by Sullivan et al. (2017). Whether the role of viruses in C cycling is more defined by ^“^viral shunt^”^ or ^“^viral shuttle^”^ when lysing host cells and how it may differ in other ecosystems such as soils, remains undetermined.

Soil systems possess hierarchical pore structure with associated pore accessibility for community members being influenced by the hydrologic landscape (Xu et al., 2017; Kuzyakov & Mason-Jones, 2018). Differences between soil and aquatic ecosystems could shape distinct virus-host interaction patterns, and the mechanism of viruses impacting host community and large-scale environmental processes may also differ. Recently, new data to test the existing paradigms of viral impact on C and nutrient cycling in soil has begun to emerge (Albright et al., 2022; Wang et al., 2022). Direct experimental efforts to examine the impact of soil viruses on ecosystem processes are still scarce. Viruses affect C flow not only by influencing microbial metabolism but also via the fate of residual C resulting from viral-mediated cell lysis. Here, we conducted microcosm experiments by manipulating virus and cell abundances by adding extracted soil viruses and bacteria back to sterile sand or soil. The added viruses and bacteria were derived from either agricultural (AG) or grass land meadow steppe (GL) soil. The viral impact was assessed by comparing the virus treatment (microcosms with active soil viruses) with the control (microcosms with inactivated soil viruses). We monitored variations in virus and bacteria abundance, bacterial community structure, organic C mineralization (i.e., respiration), nitrous oxide (N_2_O) emissions, and soil organic C content in the microcosms over 24 days. This study aims to reveal the critical role of soil viruses in regulating microbial community succession and biomass turnover and interrogate the biogeochemical consequences.

## 2. Materials and methods

### 2.1. Site description, soil sampling and processing

Soils were sampled from an agricultural site of maize monoculture (AG) at Shenyang Ecological Experimental Station (41°31^’^ N, 123°24^’^ E) and from a meadow steppe (GL) at Erguna Forest-Steppe Ecotone Research Station (50°12′N,119°30′E). Shenyang Ecological Experimental Station sits in the center of the southern Songliao Plain, with an average altitude of 41 m. It has a temperate subhumid continental monsoon climate with four distinct seasons, and the average annual temperature is 7-8 ℃. The soil sampling site of Erguna research station lies in Heishantou Town, Erguna City, Hulunbuir City, Inner Mongolia Autonomous Region, and is 523 m above sea level. This region is subject to the temperate and cold temperate climate, with short frost-free period, hot and dry summer, and long winter.

Surface soil samples (0-10 cm) were randomly taken from five locations at each site by using coring devices. The soil samples from each site were combined and thoroughly mixed. Each of the soil samples was placed into a plastic bag and appropriately labeled. All samples were immediately transported back to the laboratory on ice for further analyses. The coring devices were cleaned and sterilized with ethanol (70%, v/v) before sampling to prevent contamination.

All samples were sieved with a 2-mm sieve. Before microcosm experiments, 80 ml of minimal medium supplemented with glucose was added to 300 g of sieved soil which was then thoroughly mixed. The soil samples were incubated at 25 ℃, for 48h to enrich the soil microbial and viral communities. Minimal medium contained 1.265 g L^-1^ K_2_HPO_4_, 1.125 g L^-1^ Na_2_HPO_4_, 0.25 g L^-1^ (NH_4_)_2_SO_4_, 0.049 g L^-1^ MgSO_4_ and 2.702 g L^-1^ C_6_H_12_O_6_. After incubation, soil bacteria and extracellular viruses were extracted for subsequent inoculation in the experimental sand microcosm system.

### 2.2. Extraction and enumeration of soil microbes and viruses

Extraction of viruses and microbes from soils was performed as described in Williamson et al. (2003) and illustrated in Fig. S1. The cultured soils were suspended with cold extraction buffer (containing 1.44 g L^-1^ Na_2_HPO_4_⋅7H_2_O, 0.24 g L^-1^ KH_2_PO_4_, and 10 g L^-1^ potassium citrate, pH 7) in a w/v ratio of 3:10 (Williamson et al., 2003). The mixture was blended at the highest speed for 5 min. For separation of microbial cells, 30 mL of soil slurries were gently pipetted onto 6 ml 60% (w/v) nycodenz solution in a 50 mL centrifuge tube and were centrifuged at 5000 × g for 30 min. The supernatant including the nycodenz phase was collected to ensure acquisition of the most extractable bacterial cells. The harvested bacterial extracts (the supernatant including the nycodenz phase) from each type of soil sample, i.e., AG and GL were pooled. The microbial cells from each soil sample were captured by filtering the whole fractions of each bacterial extract onto a sterile 0.22-µm-pore-size Corning bottle top vacuum filter. The captured bacterial cells were resuspended with the same volume of sterilized minimal medium.

For separation of soil viruses, the vigorously blended soil slurries were centrifuged at 5000 × g for 30 min, and the supernatants were collected. The filtrates (viral extracts) obtained from each soil type (AG and GL) were converged and filtered through a sterile 0.22-µm-pore-size Corning bottle top vacuum filter to remove cellular-size particles. The viral extractions were purified and concentrated by using Amicon Ultra-15 centrifugal filters (15 mL, 30 kDA, Millipore, Massachusetts). Half of the viral extracts were inactivated by autoclaving at 121℃ for 20 min, thrice. All the microbial and viral extracts were immediately used for microcosm experiments.

Microbial and virus-like particles (VLPs) abundances were determined through epifluorescence microscopy enumeration as previously described (Williamson et al., 2003; Liang et al. 2020). Before enumeration, the unencapsulated DNA in the viral and bacterial extracts were digested with deoxyribonuclease (DNase) I enzyme. After DNase treatment, appropriate volume (500 μL to 1 mL, based on the density of VLPs) of viral extract from each sample was vacuum filtered through a 0.02-μm-pore-size Whatman Anodisc filter, while each bacterial extract was vacuum filtered through a 0.2-μm Isopore membrane filter (Merck Millipore Ltd., Cork, Ireland). The VLPs and bacteria were trapped on the corresponding filters, which were thereafter stained with 2X SYBR gold nucleic acid dye (1:5000 dilution of original stock, Invitrogen). The prepared filters were immediately observed at 1000X magnification using an Olympus BX53 epifluorescence microscopy (filter set of FITC: excitation at 467–498 nm wavelength and emission at 513–556 nm). Ten to fifteen images were taken for each sample and analyzed by using automatic counting software.

### 2.3. Microcosm experiments

The viral impact on microbial community structure and C cycling was investigated in two types of soils, i.e., an agricultural soil (AG) and a meadow steppe soil (GL). The microcosm experiment design is shown in Fig. 1. The microcosm experiment was performed in 500 mL glass media bottles, each with a cap equipped with a three-way valve for gas collection. For each microcosm, microbial extract (from 30 g of soil), minimal medium (25 mL), and viral extract (from 30 g of soil, active or inactivated) were added to 300 g sterilized sand or soil (Albright et al., 2022), which were then evenly mixed before being transferred into a sterilized glass media bottle. Specifically, the extracted bacteria and viruses from 1 g of soil samples were added into 10 g of the corresponding sterilized sand or soil (3.7 × 10^6^ bacteria and 2.3 × 10^7^ per g sand/soil for GL microcosms; 5.5 × 10^5^ bacteria and 9.6 × 10^6^ per g sand/soil for AG microcosms).

**Fig. 1.**
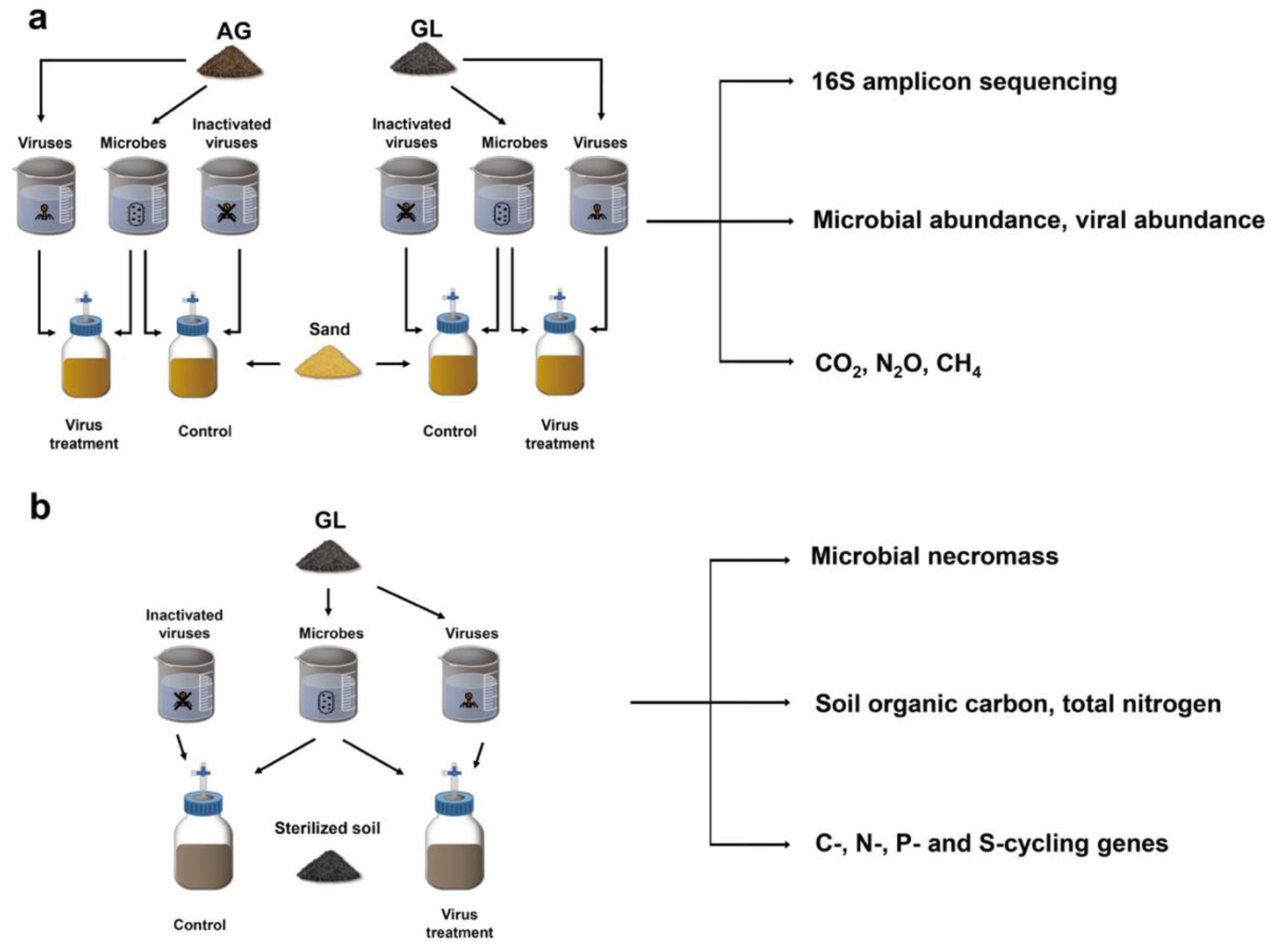
Experimental overview describing microcosm experiment design. Soil microbes and viruses were extracted from an agricultural soil of maize monoculture field (AG) and a meadow steppe soil (GL). In sand microcosms, bacterial extract (from 30 g of soil), minimal medium (25 mL), and viral extract (from 30 g of soil, active or inactivated) were added to 300 g sterilized sand (a) or soil (b). Four replicates were included in each treatment. Sand and soil microcosms were destructively sampled at 2, 5 and 10 days. Microbial abundance, viral abundance, and bacterial community structure in the sand samples were assessed. The production of CO_2_ and N_2_O in sand microcosms was monitored all through the incubation. Bacterial, fungal, and total microbial necromass C content, along with SOC and TN content, were determined in the soil microcosm samples.

The sand/soil microcosms with active viruses were the virus treatment, while the microcosms with inactivated soil viruses served as the control group. Each treatment had four replicates. Microcosms were incubated at 25℃, and gaseous production, bacterial abundance and community structure, and viral abundance were monitored throughout the incubation period. Fluxes of greenhouse gasses (i.e., CO_2_, CH_4_, and N_2_O) were determined at 0.5, 1, 2, 3, 4, 5, 6, 7, 9, 14, 24, and 25 d. Soil microcosms were destructively sampled at 2, 5, and 10d, and the soil organic C (SOC), total nitrogen (TN), and microbial necromass content (calculated using the content of amino sugars) in each soil microcosm was determined.

### 2.4. Amplicon sequencing and analysis of bacterial community structure

The total soil DNA in each sample was extracted by using PowerLyser PowerSoil DNA isolation kit (Qiagen, Hilden, Germany). The quantification of the DNA samples was performed by using Quant-iT PicoGreen dsDNA assay kit (Invitrogen, Carlsbad, CA, US). The qualified DNA samples were sent to Guangdong Magigene Biotechnology Co. Ltd. for Next-Generation sequencing of the V3 and V4 regions of 16S rRNA genes. PCR amplifications were employed for preparation of the V3–V4 gene libraries, and the amplicon sequencing was performed on an Illumina Nova6000 platform. The 16S rRNA gene sequencing data was processed with QIIME2 DADA2 pipeline (version 1.16) (Callahan et al., 2016) for evaluating the bacterial community taxonomic profiles. In brief, the raw sequences were quality-filtered, trimmed, and merged using VSEARCH (version 1.9.6) and CUTADAPT (version 1.9.1). The obtained high-quality amplicon sequences were aligned to a customized reference alignment of SILVA-based bacterial reference (version of v138) and were clustered into operational taxonomic units (OTUs) according to UPARSE algorithm (sequence identity similarity threshold at 97%). The rarefaction curves showed that the bacterial species richness tended to approach the saturation plateau in all samples, suggesting sufficient sequencing depth for describing the bacterial community diversity. Shannon index in each sample was calculated to estimate the bacterial community diversity. The bacterial community evenness was also analyzed. Principal co-ordinates analysis (PCoA) was performed to show the dissimilarities among the bacterial communities. The sequence data was deposited at NCBI database and can be accessed under project PRJNA1033447.

### 2.5. Analysis of microbial functional genes

The high-throughput quantitative PCR (qPCR)-based chip was used for determining the abundance of microbial functional genes involved in C, N, phosphorus (P) and sulfur (S) cycles (Zheng et al., 2018). Briefly, the quantity and quality of the extracted total soil DNA from each sample were assessed on NanoDrop ND-2000 (ThermoScientific, Wilmington, DE, USA). DNA samples, qPCR reagents, and qPCR primers for 71 microbial C-, N-, P- and S-cycling genes were added onto a nested nanowell high-throughput qPCR chip. The qPCR reactions and fluorescence detection were performed on a SmartChip Real-Time PCR System. The abundance of 16S rRNA genes was used as the reference metric for absolute quantification of all genes.

### 2.6. CO_2_, CH_4_, and N_2_O emission

On days of 0.5, 1, 2, 3, 4, 5, 6, 7, 9, 14, 24, and 25, gas samples (35 mL) were collected from each microcosm and stored in glass vials. The CO_2_, CH_4_, and N_2_O emissions were analyzed simultaneously with an Agilent 7890A gas chromatograph (Wilmington, DE, USA) equipped with flame ionization (FID) and electron capture detectors (ECD) (Zhang et al., 2022). The relative concentrations of each gas were calculated as the following equation:

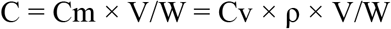

where C (mg/kg) is the relative concentration of CO_2_, CH_4_, and N_2_O; Cm (mg/L) and Cv (mL/L) are the mass and volume concentration of the gaseous products, respectively; V (L) is the volume of head space; ρ (mg/L) is gas density under standard conditions; W (kg) is the soil dry weight. The headspace air in each microcosm was replaced by sterilized air after each measurement.

### 2.7. Soil properties

Sand/soil samples were destructively collected from microcosms on the 2^nd^, 5^th^, and 10^th^ days of incubation. The soil water content was measured by drying the fresh soil in an oven. For determining the content of the SOC, TN, and amino sugars in soil, the soil samples were air dried and ground before the analysis. The SOC and TN content in each sample were measured using a Flash Smart Elemental Analyzer (Thermo Fisher Scientific). Amino sugars, used as indices for soil microbial necromass accrual, were determined as previously described by Zhang and Amelung (1996). Specifically, three amino sugars, i.e., muramic acid (MurA), glucosamine (GluN), and galactosamine (GalN), were measured for quantification of microbial necromass accumulation in each soil sample. The bacterial and fungal necromass C contents were calculated according to the following equations derived from Liang et al. (2019). Bacterial necromass C = MurA × 45; fungal necromass C = (GluN/179.17 - 2 × MurA/251.23) × 179.17 × 9. The sum of bacterial and fungal necromass C was used to assess the contribution of the total microbial necromass C to SOC in each soil sample.

### 2.8. Statistical analysis

Statistical analyses of bacterial community composition were performed in the R software environment. Principal coordinates analysis (PCoA) based on UniFrac analyses was used to compare the bacterial communities in all samples. A mixed model with repeated measures was performed for the cumulative CO_2_ and N_2_O release, bacterial taxonomic composition, functional gene composition, SOC, TN, and microbial necromass C. Virus treatment and soil types were fixed effects, with time series being regarded as repeated measures and the replicates as random effects. The two-way analysis of variance (ANOVA) followed by Duncan^’^s post-hoc test was used to examine the effects of virus treatment and soil types on CO_2_, CH_4_, and N_2_O production, bacterial taxonomic composition, and soil physiochemical properties, with statistical analyses being performed with IBM SPSS 19.0 software package (IBM Corporation, Chicago, USA). The major results were visualized in the R software environment and Microsoft Excel 2020. The plausible interaction paths between viral addition, microbial community traits, greenhouse gas emissions and SOC sequestration were assessed via Pearson correlation analysis on Origin2023b software. The structural equation modeling (SEM) was performed in AMOS software (17.0.2 student version; Amos Development, Crawfordville, FL, USA) to establish direct and indirect relationships between SOC mineralization and tested parameters. By increasing or decreasing variables in the SEM, a pathway model that can optimally indicate the effects of soil viruses on microbial community dynamics, soil nutrients turnover, and SOC sequestration was gradually obtained.

## 3. Results

### 3.1. The impact of soil viruses on microbial community succession

The 16S rRNA gene amplicon sequencing showed that co-inoculation of soils with native bacteria and viruses significantly altered the bacterial community composition of the agricultural soil but not the grassland soil (Fig. 2a). In the AG sand microcosms, the virus treatment samples were clearly separated from the control samples (Fig. 2b). Viral impacts substantially altered the relative abundances of the dominant AG soil bacterial taxa added to sand. Specifically, virus treatment sharply decreased the proportions of *Pseudomonas, Acinetobacter, Flavobacterium, Pedobacter, Chryseomicrobium, Exiguobacterium, Stenotrophomonas, Rhizobiaceae*, and *Bacillus*, while *Comamonas, Brevundimonas, Erwiniaceae*, and *Enterobacteriaceae* were dramatically enriched (Fig. 2c). In contrast, the succession of GL bacterial communities was not significantly altered by soil viruses, as the incubation time was the primary factor influencing bacterial community composition (Fig. 2d). The bacterial community structure in the GL microcosms clustered by incubation time rather than virus treatment. Though the beta diversity clearly varied with incubation time, the effect of viral addition on bacterial taxonomic composition of GL sand-microcosms was similar to that in AG sand-microcosms, but the extent to which viruses increased or decreased the relative abundances of specific taxonomic groups was much lower. Soil viruses significantly increased bacterial diversity in AG sand-microcosms at day 5 (based on Shannon and Evenness indices, *P* < 0.05), however the bacterial diversity in GL sand-microcosms was not significantly influenced by viral addition (Fig. 2e). Soil viruses increased bacterial community diversity and evenness at day 5, but there was no significant change in bacterial community diversity along the incubation period (Fig. 2e).

**Fig. 2.**
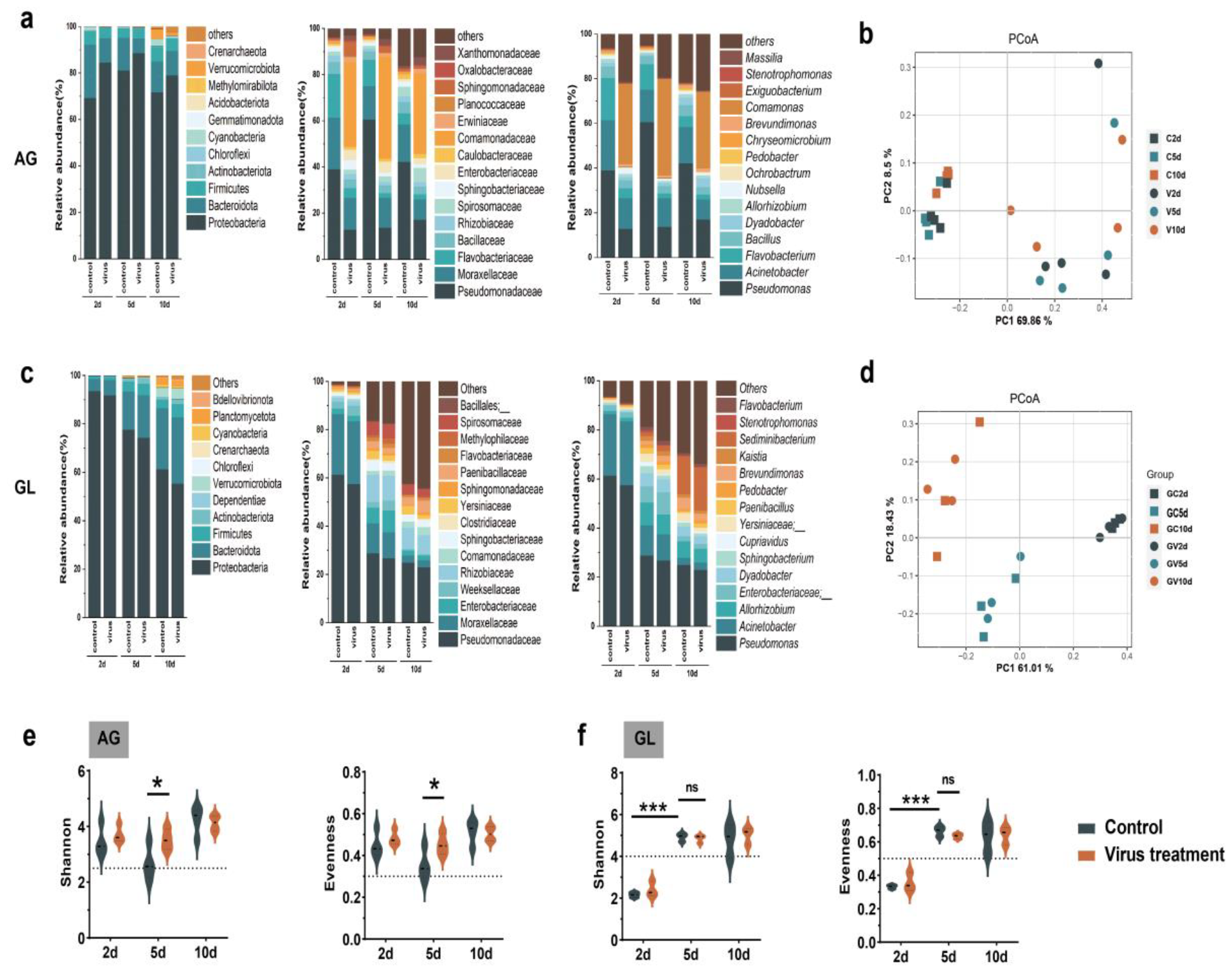
The impacts of soil viruses on bacterial community dynamics in AG and GL sand-microcosms. The taxonomic composition of bacterial community in AG (a) and GL (b) microcosms was explored at phylum-, class- and genus-level. Principal Coordinate Analysis (PCoA) was performed to estimate the dissimilarities among the bacterial communities in different samples of AG (c) and GL (d) microcosms. The bacterial community Shannon diversity index and evenness in AG and GL microcosms were shown in e and f, respectively.

The abundance of virus-like particles (VLPs) ranged from 7.1 × 10^8^ to 7.6 × 10^8^ in AG microcosms and 3.9 × 10^8^ to 5.1 × 10^8^ in GL sand-microcosms (Fig. 3a). Interestingly, the bacterial abundance increased in soils of the virus treatment compared to that of control microcosms (*P* < 0.01 for AG and *P* > 0.05 for GL; Fig. 3b), suggesting viruses may increase the experimental system^’^s capacity of sustaining bacterial populations presumably by restructuring cellular metabolic network and altering community turnover rate (Forterre, 2013; Howard-Varona et al. 2020).

**Fig. 3.**
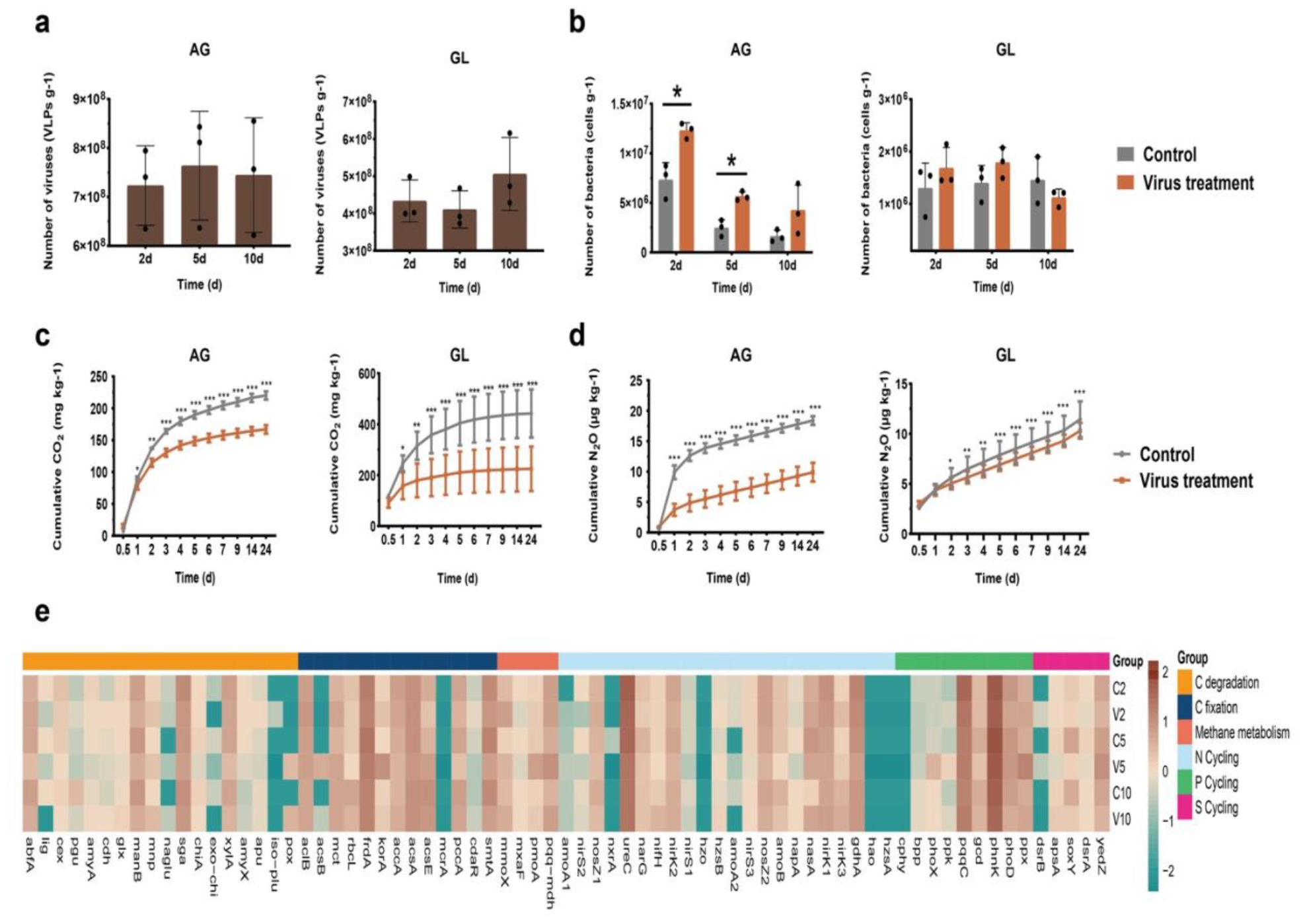
The viral abundance (a), bacterial abundance (b), and production of greenhouse gases, i.e., CO_2_ (c) and N_2_O (d), in AG and GL sand-microcosms. The profiles of C-, N-, P- and S-cycling genes were shown in e. Each data point represents the mean value of three replicate samples. The error bars represent standard deviation. Statistical significance between virus treatment and the control group is indicated by ^**\***^ *P* < 0.05, ^**\*\***^ *P* < 0.01, and ^**\*\*\***^ *P* < 0.001.

### 3.2. Greenhouse gas production and microbial functional gene profile

Bacterial metabolism transforms C and N sources into bacterial biomass while releasing gaseous products, e.g., CO_2_ and N_2_O, into the environment (Jansson and Hofmockel, 2020). Virus addition significantly reduced CO_2_ production in both AG (25% reduction compared to control) and GL (91% reduction) sand-microcosms (Fig. 3c). Soil virus addition also reduced N_2_O production in AG but not the GL sand-microcosms by 45% leading to less gaseous N loss from the experimental system (Fig. 3d). The production of CH_4_ was not influenced by soil viruses in both AG and GL sand-microcosms (Fig. S2). These results suggest that viral activity likely plays a role in net emissions of this potent greenhouse gas.

Distinct microbial functional profiles of C, N, P and S cycling were observed across different samples (Fig. 3e). In the early stage of the experiment, viral addition sharply decreased the abundances of the assessed functional genes (a reduction of 50% at day 2; Fig. S3). Conversely, the quantities of the C-, N-, P- and S-cycling genes were significantly increased at day 5 and 10. Specifically, *glx* (C degradation), *mnp* (C degradation), *amyX* (C degradation), *apu* (C degradation), *aclB* (C fixation), *mct* (C fixation), *frdA* (C fixation), *korA* (C fixation), *accA* (C fixation), *acsA* (C fixation), *mmoX* (methane metabolism), *pmoA* (methane metabolism), *pqq-mdh* (methane metabolism), *ureC* (N cycling), *narG* (N cycling), *nifH* (N cycling), *nirK2* (N cycling), *hzsB* (N cycling), *nirS3* (N cycling), *nosZ2* (N cycling), *nirK1* (N cycling), *nirK3* (N cycling), *pqqC* (P cycling), *gcd* (P cycling), *phnK* (P cycling), *ppx* (P cycling), *dsrA* (S cycling) and *apsA* (S cycling) were enriched in the soils of virus treatment (Fig. 3e). Soil viruses also reduced the abundances of *cex* (C degradation), *amyA* (C degradation), *chiA* (C degradation), *xylA* (C degradation), *pccA* (C fixation), *smtA* (C fixation), *nirS2* (N cycling), *amoB* (N cycling), *napA* (N cycling), and *soxY* (S cycling). Soil viruses decreased microbial functional diversity (based on Shannon index) at day 2 while increasing the Shannon diversity at day 5 and 10.

### 3.3. Microbial necromass accumulation and its relationship with SOC

In GL soil-microcosms, the effects of viral addition on soil organic C transformation and accrual were explored, showing that soil viruses significantly increased the content of the three measured amino sugars (*i*.*e*., MurA, GlcN, and GalN, important indices for the contribution of microbial residues to soil organic matter), in soils across all microcosms (*P* < 0.05; Fig. S4). The total microbial necromass C content in the virus treatment microcosms remarkably increased by 4.3-8.3% (bacterial necromass C: 10.1-13.1%; fungal necromass C: 2.2-7%) compared with that in the control (*P* < 0.05; Fig. 4a). Soil properties analysis showed that soil viruses slightly increased SOC and TN content, especially during the late stage of the microcosm experiment, i.e., at day 10 (*P* < 0.05) and 45(*P* < 0.05) (Fig. 4b, c). The markedly lowered organic C mineralization rate and increased microbial abundance, combined with promoted microbial biomass accrual, suggested that viral infection caused higher microbial CUE. Correlation analysis suggests that SOC content was negatively correlated with CO_2_ production (*r* = -0.75, *P* < 0.001; Pearson correlation) but positively correlated with microbial necromass C accrual (*r* = 0.47, *P* < 0.05; Pearson correlation) (Fig. S3), both of which were tightly related to viral infections.

**Fig. 4.**
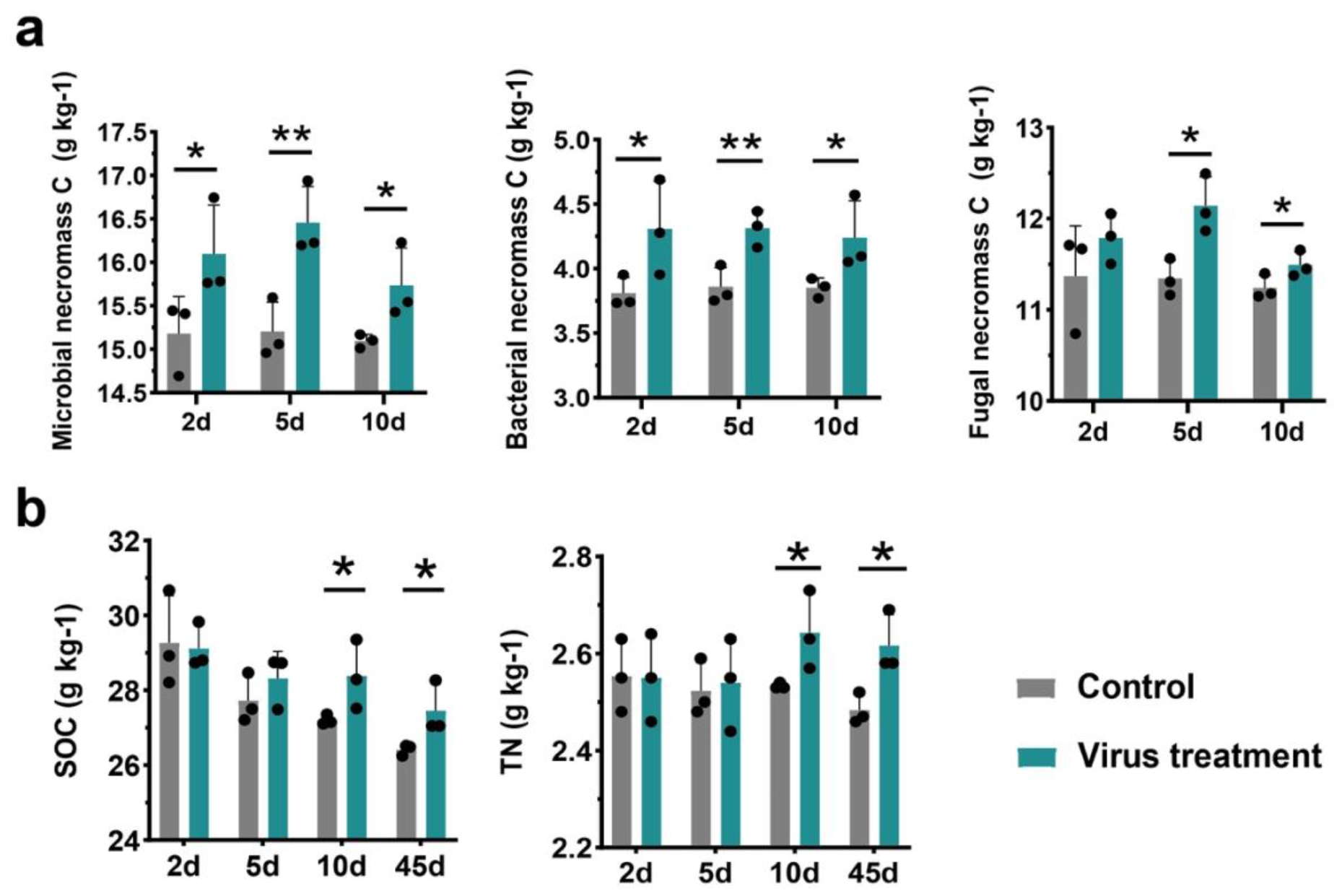
Microbial necromass C (a), soil organic C (b), and total nitrogen (c) content in soil microcosms. Bacterial necromass C was calculated based on the content of muramic acid (MurA), and fungal necromass C was derived from content of MurA and glucosamine (GluN) according to the previously described equation (Liang et al., 2019). The total microbial necromass C was the sum of bacterial and fungal necromass C.

### 3.4. Linking viral infections to SOC accumulation

The reduced carbon mineralization and accelerated microbial necromass accumulation suggest that viral infections directly influence SOC transformation and consequently may affect C sequestration. Structural equation modeling (SEM), a hypothesis-driven multivariate data analysis technique, was performed to assess how viral infections were related to microbial community properties and C sequestration in soil (Fig. 5a). Soil viruses significantly altered microbial abundance, community diversity and C-cycling gene profiles (Fig. S5). Viral infections can also control CO_2_ production and contribute to bacterial necromass accrual. Bacterial necromass C positively affected C sequestration by increasing the contribution of microbial necromass to total SOC. Soil viruses were also positively related to SOC content via CO_2_ emission reduction.

**Fig. 5.**
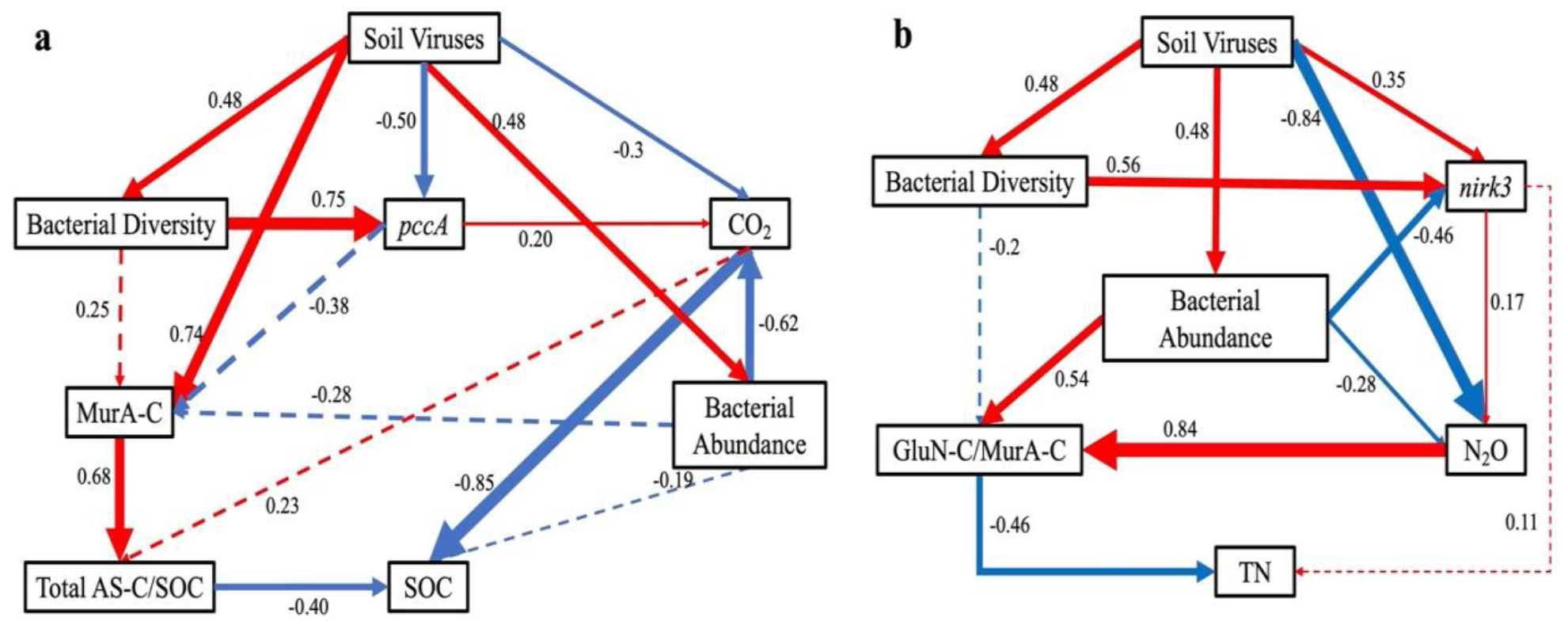
Structural equation model relating soil viruses to microbial community dynamics, soil nutrients turnover, TN (a), and SOC sequestration (b). Solid arrows denote direct effects of a variable on another, with red color representing positive relationships and blue corresponding to negative ones (*P* < 0.05). Arrow thickness and the number represent the magnitude of the standardized path coefficient. Non-significant pathways in the model are shown with dashed arrows. MurA-C: bacterial necromass C; Total AS-C/SOC: the contribution of total microbial necromass C to SOC; GluN-C/MurA-C: the ratio of fungal necromass C to bacterial necromass C.

Furthermore, the relationship between viral infections and TN content in soil was examined in SEM. Soil viruses positively affected soil TN content through impacts on microbial abundance, N-cycling gene composition, N_2_O production, and fungal:bacterial necromass ratio (Fig. 5b). The microbial diversity, abundance, functionality, and production of greenhouse gases appeared to be the crucial pathways in both structural equation models to link viral infections to enhanced SOC and TN content.

## 4. Discussion

### 4.1 Microbial community dynamics affected by soil viruses

Viral infections have been shown as a major driver of microbial mortality and are tightly related to microbial community diversity. For example, according to the widely accepted ^“^kill-the-winner^”^ model, the more abundant microbial populations are more susceptible to viral lysis, therefore increasing microbial community evenness and diversity (Chevallereau et al., 2022; Liang et al., 2023). In this study, viral addition clearly impacted microbial community structure, but the microbial communities in AG and GL sand-microcosms were impacted differently, with the microbial community structure in AG sand-microcosms being more substantially restructured by viral infections. The causes of these differences are unknown but involve the nature of the soil ecosystems from which the viruses and bacteria used as inoculum were obtained. Further research is needed to elucidate the cause(s) for these observations. The prominently different effects of soil viruses on microbial community structure may be explained as follow: (1) The microbial populations in AG and GL soil microcosms were clearly different, thus possibly having contrasting virus-host interaction patterns. (2) The unique soil physiochemical properties in AG and GL soils may have cultivated stable relationships between soil viruses and their hosts. (3) Moreover, compared with the undisturbed GL soils, the AG soil was subject to consistent anthropogenic-management practices, therefore the microbial and viral communities from AG soil were more sensitive to transient changes. Indeed, the GL soil contained two-fold more SOC (GL: 23.9 g kg^-^ _1_; AG: 12.3 g kg^-1^) and TN (GL: 2.2 g kg^-1^; AG: 1.1 g kg^-1^) content than the AG soil (Fig. S6). The microbial necromass content in GL soil (14.1 g kg^-1^) was nearly three-fold higher than that in AG soil (4.9 g kg^-1^) (Fig. S6). The GL soil was more nutrient-rich supporting higher microbial growth rates, and according to ^“^piggyback-the-winner^”^ paradigm, viral lysis may have a less important role in shaping microbial community structure (Silveira et al., 2021). However, the effects of soil viruses on the succession of microbial community in AG soil were in agreement with ^“^kill-the-winner^”^ model, as the relative abundances of the dominant bacterial populations, such as *Pseudomonas, Acinetobacter, Flavobacterium*, and *Bacillus*, were dramatically reduced by viral infections, leading to increases in other bacterial populations (*e*.*g*., *Comamonas* and *Brevundimonas*). These bacterial taxa, frequently observed as key nodes in microbial interaction networks, were also shown to be evidently affected by viral infections in other studies (Braga et al., 2020; Liu et al., 2023). Therefore, soil viruses can influence microbial community succession not only by infecting the abundant groups but also by restructuring the interactions within the community.

### 4.2 Soil viruses modulate C and N cycling

By assessing the impacts of soil viruses on microbial community succession and function, we demonstrated that soil viruses can influence microbial CUE and community turnover rate, causing significant reduction in GHGs emissions. The microbial communities responded quite differently in the AG and GL soil microcosms. Interestingly, though the bacterial community responses to virus addition were different, the functional response with respect to respiration were similar yet with no significant inhibition of N_2_O emission in response to virus addition observed for the GL microcosms. Presumably the organic N released from bacteria lysed by viral infection was mineralized to NH_4_^+^, but nitrification was apparently not inhibited in the virus-amended GL microcosms. The potential impact of soil viruses on N cycling has been explored in recent research. For example, Braga et al. (2020) showed that soil viruses can influence N availability in soil with ammonium positively related to the presence of viruses. Moreover, viral infections of nitrifiers, i.e., ammonia-oxidising archaea, were found as a frequent process in nitrification with potential consequences in microbial activity and N cycling (Lee et al., 2023).

Soil microorganisms, as decomposers and contributors to recalcitrant organic matter components, have major roles in SOC transformation and C sequestration (Liang et al., 2017; Buckeridge et al., 2022; Wang et al., 2023). Mineralization of SOC to CO_2_ by microorganisms (decomposing ∼ 60 Pg SOC per year) is a pivotal source of greenhouse gas to atmosphere, thereby being tightly linked to C storage in soil and global climate change (Hursh et al., 2017; García-Palacios et al., 2021). Microbial production and the microbially derived necromass C, *e*.*g*., dead microbial cells, cellular debris, extracellular macromolecules, and their degradants, can account for more than half of SOC in many systems (Liang et al., 2019; Wang et al., 2021; Whalen et al., 2022). The major contribution of microbial residues to SOC indicates that microbial necromass can persist and accumulate in soil over time. However, the formation of necromass-derived SOC and its subsequent fate in soil remains elusive. Recent studies showed that the death pathways of soil microbes substantially affect microbial necromass characteristics (Camenzind et al., 2023). As viral reproduction can deplete N and P within host cells, the viral lysis-caused microbial necromass was more resistant to decomposition.

Viruses can exploit host machinery and actively reprogram host metabolic networks for viral reproduction (Howard-Varona et al., 2020). Our results showed that soil viruses not only increased living microbial biomass but also stimulated microbial necromass accumulation, suggesting viral infections caused higher microbial CUE and community turnover rate. Therefore, viral infections renovated organic C partitioning by soil microbial community allocating more C towards microbial biomass and by-products. In our study, viral infections substantially influenced microbial C-, N-, P- and S-cycling genes, which may partially explain the increased microbial CUE. A recent global-scale meta-analysis examining the relationship between microbial CUE and SOC preservation demonstrated the central roles of microbial CUE in promoting SOC storage across different soil ecosystems (Tao et al., 2023). The higher microbial abundance and necromass content in the soils of virus treatment indicate that microbial community turnover was accelerated under viral predation. Viral lysate-derived dissolved organic C and other labile nutrients can increase microbial metabolic efficiency and promote microbial biomass synthesis (Zhao et al., 2019; Tong et al., 2023). Accelerated microbial turnover can positively influence the contribution of microbial necromass C and N to SOC and N storage (Hagerty et al., 2014; Prommer et al., 2020). The above results, combined with SEM, show that increased microbial CUE and accelerated microbial turnover are the main mechanisms associated with increased SOC storage under viral predation.

### 4.3 Viral loop: top-down control over microbial turnover and C sequestration

Based on the above findings, we propose a conceptual framework (viral loop) of soil viruses regulating SOC transformation. Viral infections can profoundly affect microbial community turnover and are putatively the most important cause for the death of soil microorganisms. In lytic cycles of viral replication, more C and energy flow can be allocated into organic matter synthesis and minimize unnecessary energy-consuming activities via modulation of host metabolic network. Viral lysis generates large quantities of dissolved organic C and other nutrients that are susceptible to microbial reutilization and may positively affect the living conditions for soil microbes. Virus-caused mortality accelerates microbial community turnover and enhanced microbial necromass accumulation. During viral reproduction, P was disproportionately incorporated into progeny virus particles depleting most (around 87%) of the original P content from the host cells and causing imbalance of C/N/P stoichiometry in the released microbial debris (Jover et al., 2014). The overall effects of soil viruses in our experimental system reduced C mineralization and increased microbial necromass accrual and C sequestration in soil. Modeling prediction and experimental quantification of active viral infection showed that viral infections can increase per-cell release rates of extracellular C, that are major precursors of recalcitrant C forms, by 2–4 fold (Vincent et al., 2023). The increased accelerated microbial biomass turnover and decreased bioavailability of the cell lysate in viral loop can jointly contribute to increasing the SOC content and stability.

Recently, soil mineral carbon pump (MnCP) was proposed as complimentary framework for soil microbial carbon pump (MCP), to describe the critical roles of soil minerals in enhancing SOC persistence and accumulation (Xiao et al., 2023). Our study emphasized the top-down control of viral loop on MnCP and MCP, as soil viruses exert profound influences over microbial processes of SOC production and sequestration and generate an enormous amount of OC for MnCP. Collectively, the importance of viral loop in SOC sequestration needs to be acknowledged and investigated.

## 5. Conclusion

Together, the results showed soil viruses infect and cull but do not decimate host populations, directly modulating micro-food web structure and function. This study presents the first direct evidence for the virus-facilitated microbial necromass formation and accumulation. Viral infection substantially influences C and N cycling and plays a critical role in elemental sequestration in soil. We proposed ^“^viral loop^”^ to describe the top-down control of soil viruses on SOC turnover and accumulation. Moreover, some inspiring questions were raised for future research efforts. For example, it is assumed extensive proportions of bacterial populations in the experimental system were viral infected and transformed into virocells for virions production. More evidence with innovated experimental approaches is needed for quantitatively defining the ecological role of soil viruses. This study also showed that soil viruses affected bacterial community assembly differently in different soil systems. It would be important to reveal the mechanism behind the observation. Additionally, the potential link between the ecological function of soil viruses and ecosystem function and sustainability can be deduced, but how soil viruses influence soil ecosystem services, e.g., biodiversity, C storage, and biomass production, via impact on microbial communities and biogeochemical cycling remains to be elucidated.

## Acknowledgement

This work was supported by National Key R&D Program of China (award number 2022YFD1500301), Natural Science Foundation for Excellent Young Scholars of Liaoning Province, and Major Program of Institute of Applied Ecology, Chinese Academy of Sciences (award number IAEMP202201).

## References

1. Albright, M.B., Gallegos-Graves, L.V., Feeser, K.L., Montoya, K., Emerson, J.B., Shakya, M. and Dunbar, J. Experimental evidence for the impact of soil viruses on carbon cycling during surface plant litter decomposition. ISME Commun. 2, 24 (2022).

2. Bossio, D.A. et al. The role of soil carbon in natural climate solutions. Nat. Sustain. 3, 391–398 (2020).

3. Braga, L.P.P., Spor, A., Kot, W. et al. Impact of phages on soil bacterial communities and nitrogen availability under different assembly scenarios. Microbiome 8, 52 (2020).

4. Buckeridge, K.M., Mason, K.E., Ostle, N. et al. Microbial necromass carbon and nitrogen persistence are decoupled in agricultural grassland soils. Commun. Earth Environ. 3, 114 (2022).

5. Callahan, B.J., McMurdie, P.J., Rosen, M.J. et al. DADA: high-resolution sample inference from Illumina amplicon data. Nat Methods. 13, 581–583 (2016).

6. Camenzind, T., Mason-Jones, K., Mansour, I. et al. Formation of necromass-derived soil organic carbon determined by microbial death pathways. Nat. Geosci. 16, 115–122 (2023).

7. Chevallereau, A. et al. Interactions between bacterial and phage communities in natural environments. Nat. Rev. Microbiol. 20, 49–62 (2022).

8. Forterre, P. The virocell concept and environmental microbiology. ISME J. 7, 233–236 (2013).

9. Fuhrman, J.A. Marine viruses and their biogeochemical and ecological effects. Nature 399, 541–548 (1999).

10. García-Palacios, P., Crowther, T.W., Dacal, M. et al. Evidence for large microbial-mediated losses of soil carbon under anthropogenic warming. Nat. Rev. Earth Environ. 2, 507–517 (2021).

11. Hagerty, S., van Groenigen, K., Allison, S. et al. Accelerated microbial turnover but constant growth efficiency with warming in soil. Nature Clim. Change 4, 903–906 (2014).

12. Howard-Varona, C., Lindback, M.M., Bastien, G.E., et al. Phage-specific metabolic reprogramming of virocells. ISME J. 14, 881–895 (2020).

13. Hursh, A., et al. The sensitivity of soil respiration to soil temperature, moisture, and carbon supply at the global scale. Glob. Change Biol. 23, 2090–103.

14. Jansson, J.K. & Hofmockel, K.S. Soil microbiomes and climate change. Nat. Rev. Microbiol. 18, 35–46 (2020).

15. Jansson, J.K. & Wu, R. Soil viral diversity, ecology and climate change. Nat. Rev. Microbiol. (2022).

16. Jover, L., Effler, T., Buchan, A. et al. The elemental composition of virus particles: implications for marine biogeochemical cycles. Nat. Rev. Microbiol. 12, 519–528 (2014).

17. Kuzyakov, Y. & Mason-Jones, K. Viruses in soil: Nano-scale undead drivers of microbial life, biogeochemical turnover and ecosystem functions. Soil Biol. Biochem. 127, 305–317 (2018).

18. Lara, E. et al. Unveiling the role and life strategies of viruses from the surface to the dark ocean. Sci. Adv. 3, e1602565 (2017).

19. Lee, S., Sieradzki, E.T., Nicol, G.W. et al. Propagation of viral genomes by replicating ammonia-oxidising archaea during soil nitrification. ISME J. 17, 309–314 (2023)..

20. Liang, C., Schimel, J.P. & Jastrow, J.D. The importance of anabolism in microbial control over soil carbon storage. Nat. Microbiol. 2, 17105 (2017).

21. Liang, C., Amelung, W., Lehmann, J. & Kästner, M. Quantitative assessment of microbial necromass contribution to soil organic matter. Glob. Chang. Biol. 25, 3578–3590 (2019).

22. Liang, X., Radosevich, M., DeBruyn, J.M., Wilhelm, S.W., McDearis, R. & Zhuang, J. Incorporating viruses into soil ecology: a new dimension to understand biogeochemical cycling. Crit. Rev. in Env. Sci. Tec. 54, 117–137. (2024)

23. Liang, X., Wagner, R.E., Li, B., Zhang, N. & Radosevich, M. Quorum sensing signals alter in vitro soil virus abundance and bacterial community composition. Front. Microbiol. 11, 1287 (2020).

24. Liu, C., Ni, B., Wang, X., Deng, Y., Tao, L., Zhou, X. & Deng, J. Effect of forest soil viruses on bacterial community succession and the implication for soil carbon sequestration. Sci. Total Environ. 892, 164800 (2023).

25. Nicolas, A.M., Sieradzki, E.T., Pett-Ridge, J. et al. A subset of viruses thrives following microbial resuscitation during rewetting of a seasonally dry California grassland soil. Nat. Commun. 14, 5835 (2023).

26. Prommer, J. et al. Increased microbial growth, biomass, and turnover drive soil organic carbon accumulation at higher plant diversity. Glob. Change Biol. 26, 669–681 (2020).

27. Silveira, C.B., Luque, A. & Rohwer, F. The landscape of lysogeny across microbial community density, diversity and energetics. Environ. Microbiol. 23, 4098–4111 (2021).

28. Sullivan, M.B., Weitz, J.S. & Wilhelm, S. Viral ecology comes of age. Environ. Microbiol. Rep. 9, 33–35 (2017).

29. Suttle, C.A. Marine viruses—major players in the global ecosystem. Nat. Rev. Microbiol. 5, 801–812 (2007).

30. Tao, F., Huang, Y., Hungate, B.A. et al. Microbial carbon use efficiency promotes global soil carbon storage. Nature 618, 981–985 (2023).

31. Tong, D., Wang, Y., Yu, H. et al. Viral lysing can alleviate microbial nutrient limitations and accumulate recalcitrant dissolved organic matter components in soil. ISME J. 17, 1247–1256 (2023).

32. Vincent, F., Gralka, M., Schleyer, G. et al. Viral infection switches the balance between bacterial and eukaryotic recyclers of organic matter during coccolithophore blooms. Nat. Commun. 14, 510 (2023).

33. Wang, B., An, S., Liang, C., Liu, Y. & Kuzyakov, Y. Microbial necromass as the source of soil organic carbon in global ecosystems. Soil Biol. Biochem. 162, 108422 (2021).

34. Wang, C., Wang, X., Zhang, Y. et al. Integrating microbial community properties, biomass and necromass to predict cropland soil organic carbon. ISME Commun. 3, 86 (2023).

35. Wang, S. et al. Experimental evidence for the impact of phages on mineralization of soil-derived dissolved organic matter under different temperature regimes. Sci. Total Environ. 846, 157517 (2022).

36. Weinbauer, M.G. Ecology of prokaryotic viruses. FEMS Microbiol. Rev. 28, 127–181 (2004).

37. Whalen, E.D., Grandy, A.S., Sokol, N.W., et al. Clarifying the evidence for microbial- and plant-derived soil organic matter, and the path toward a more quantitative understanding. Glob. Chang. Biol. 28, 7167–85 (2021).

38. Wilhelm, S.W. & Suttle, C.A. Viruses and nutrient cycles in the sea: viruses play critical roles in the structure and function of aquatic food webs. Bioscience 49, 781–788 (1999).

39. Williamson, K.E., Wommack, K.E. & Radosevich, M. Sampling natural viral communities from soil for culture-independent analyses. Appl. Environ. Microbiol. 69, 6628–6633 (2003).

40. Xiao, KQ., Zhao, Y., Liang, C. et al. Introducing the soil mineral carbon pump. Nat. Rev. Earth Environ. 4, 135–136 (2023).

41. Xu, S., Attinti, R., Adams, E., Wei, J., Kniel, K., Zhuang, J., Jin, Y. Mutually facilitated co-transport of two different viruses through reactive porous media. Water Res. 123, 40–48. (2017)

42. Zhang, L. et al. Unexpectedly minor nitrous oxide emissions from fluvial networks draining permafrost catchments of the East Qinghai-Tibet Plateau. Nat. Commun. 13, 950 (2022).

43. Zhang, X. & Amelung, W. Gas chromatographic determination of muramic acid, glucosmine, mannosamine, and galactosamine in soils. Soil Biol. Biochem. 28, 1201–1206 (1996).

44. Zhao, Z., Gonsior, M., Schmitt-Kopplin, P. et al. Microbial transformation of virus-induced dissolved organic matter from picocyanobacteria: coupling of bacterial diversity and DOM chemodiversity. ISME J. 13, 2551–2565 (2019).

45. Zheng, B., Zhu, Y., Sardans, J. et al. QMEC: a tool for high-throughput quantitative assessment of microbial functional potential in C, N, P, and S biogeochemical cycling. Sci. China Life Sci. 61, 1451–1462 (2018).

