## Supplemental Figures for "Soil viruses reduce greenhouse gas emissions and promote microbial necromass accrual"

**Supplementary Figures**

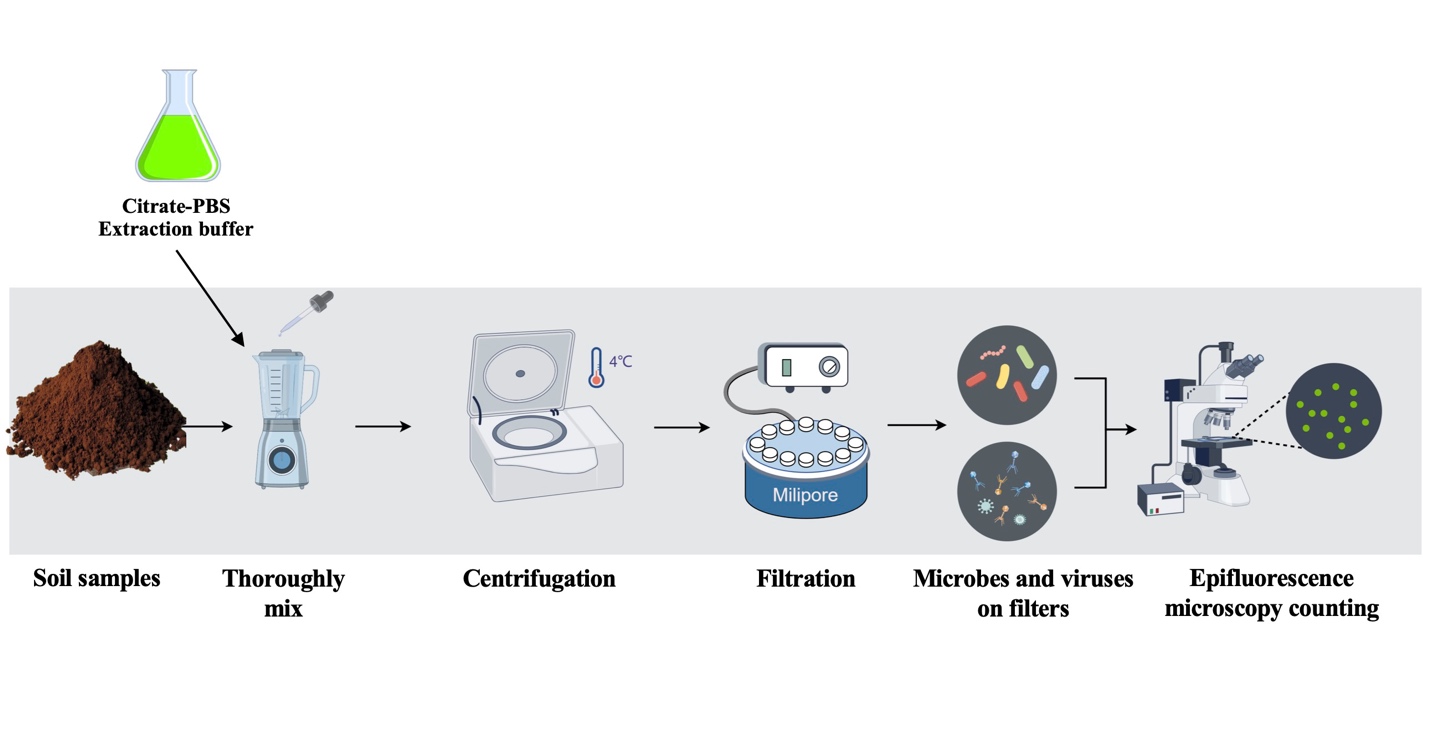
**Fig. S1** The experimental procedure of enumeration of soil microbes and viruses.

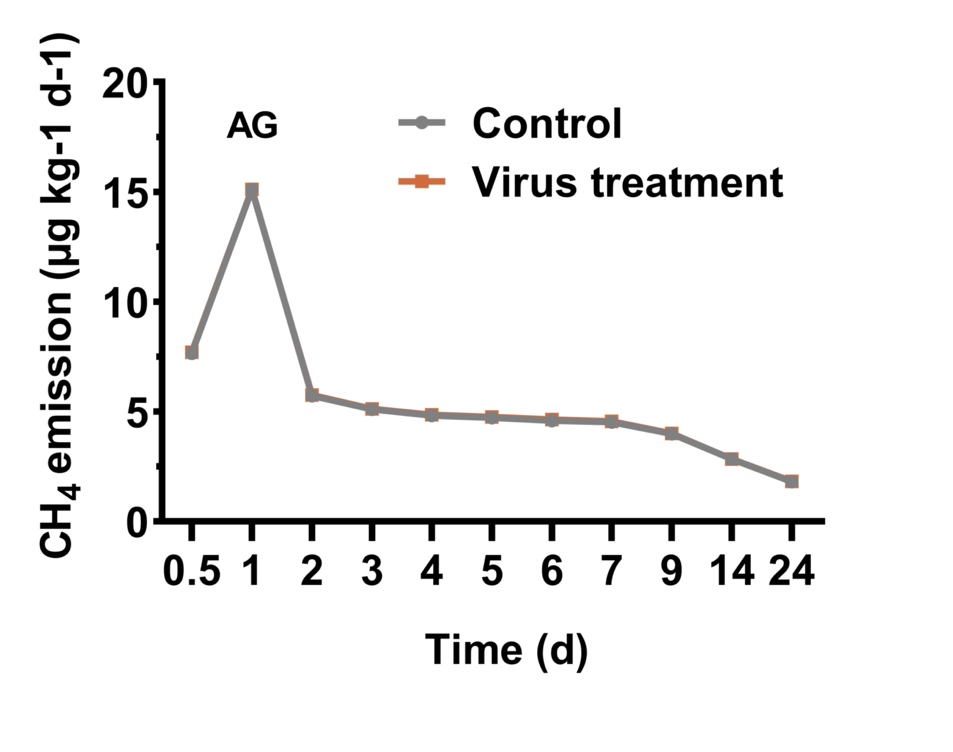

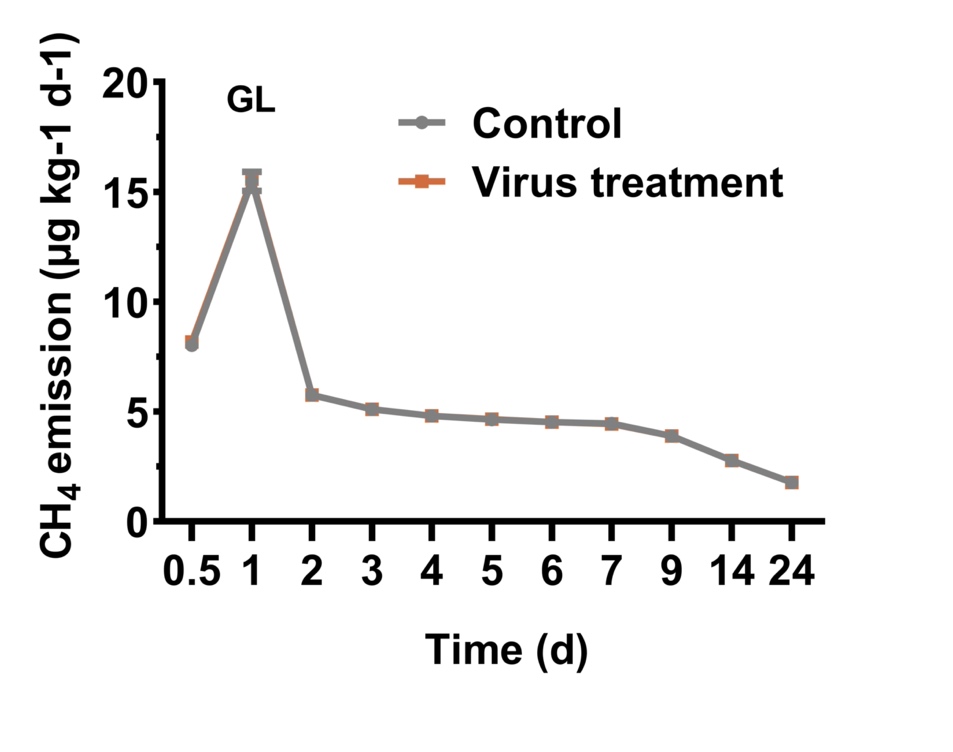

**Fig. S2** The production of CH_4_ in AG and GL sand-microcosms. Each data point represents the mean value of three replicate samples. The error bars represent standard deviation.

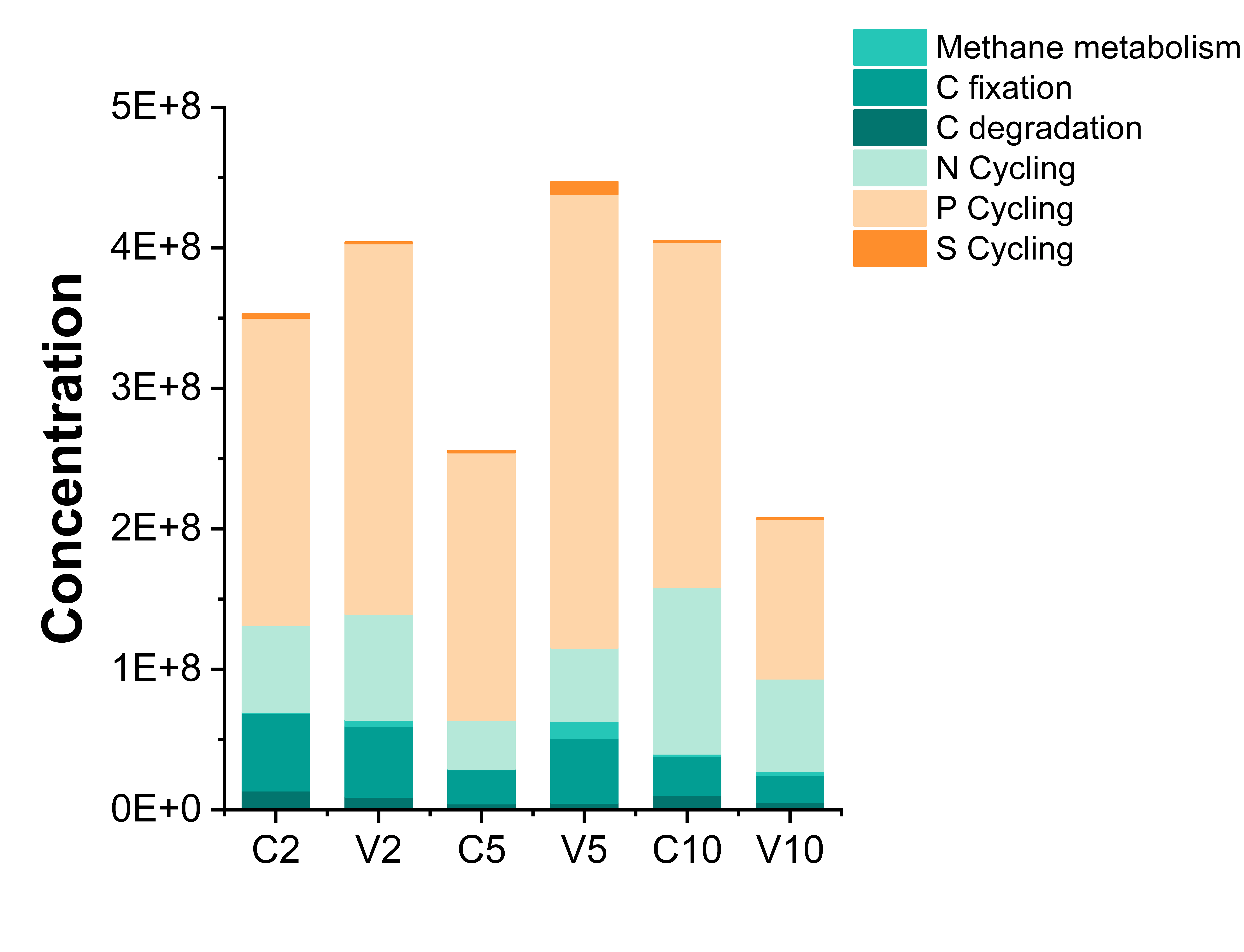

**Fig. S3** The quantities of C-, N-, P- and S-cycling genes in soil microcosms determined by high-throughput quantitative PCR (qPCR)-based chip.

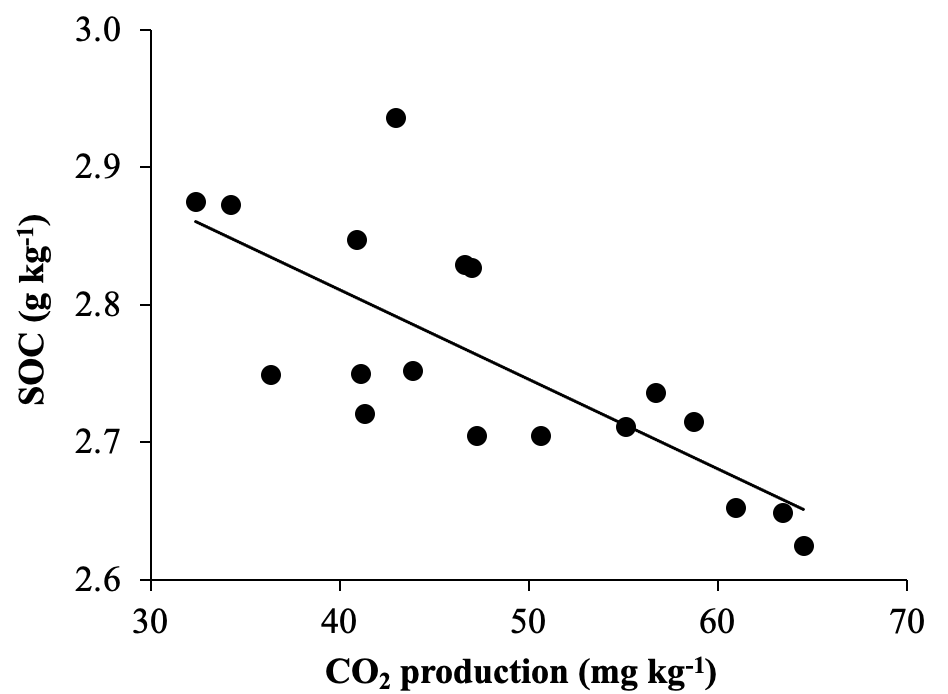

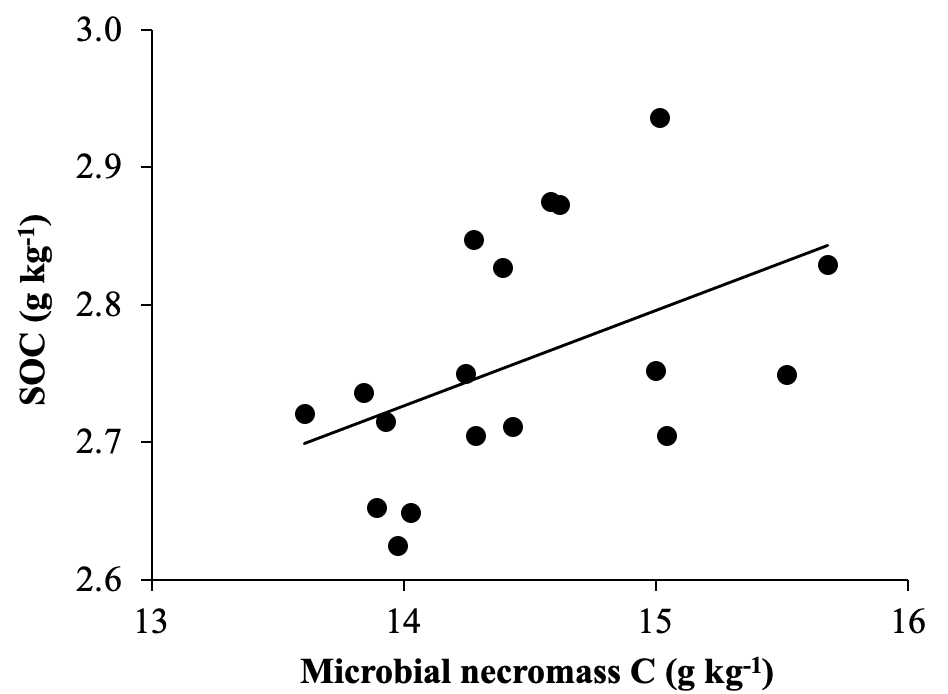

**Fig. S4** The correlation between soil organic carbon content with CO_2_ production and microbial necromass C accrual.

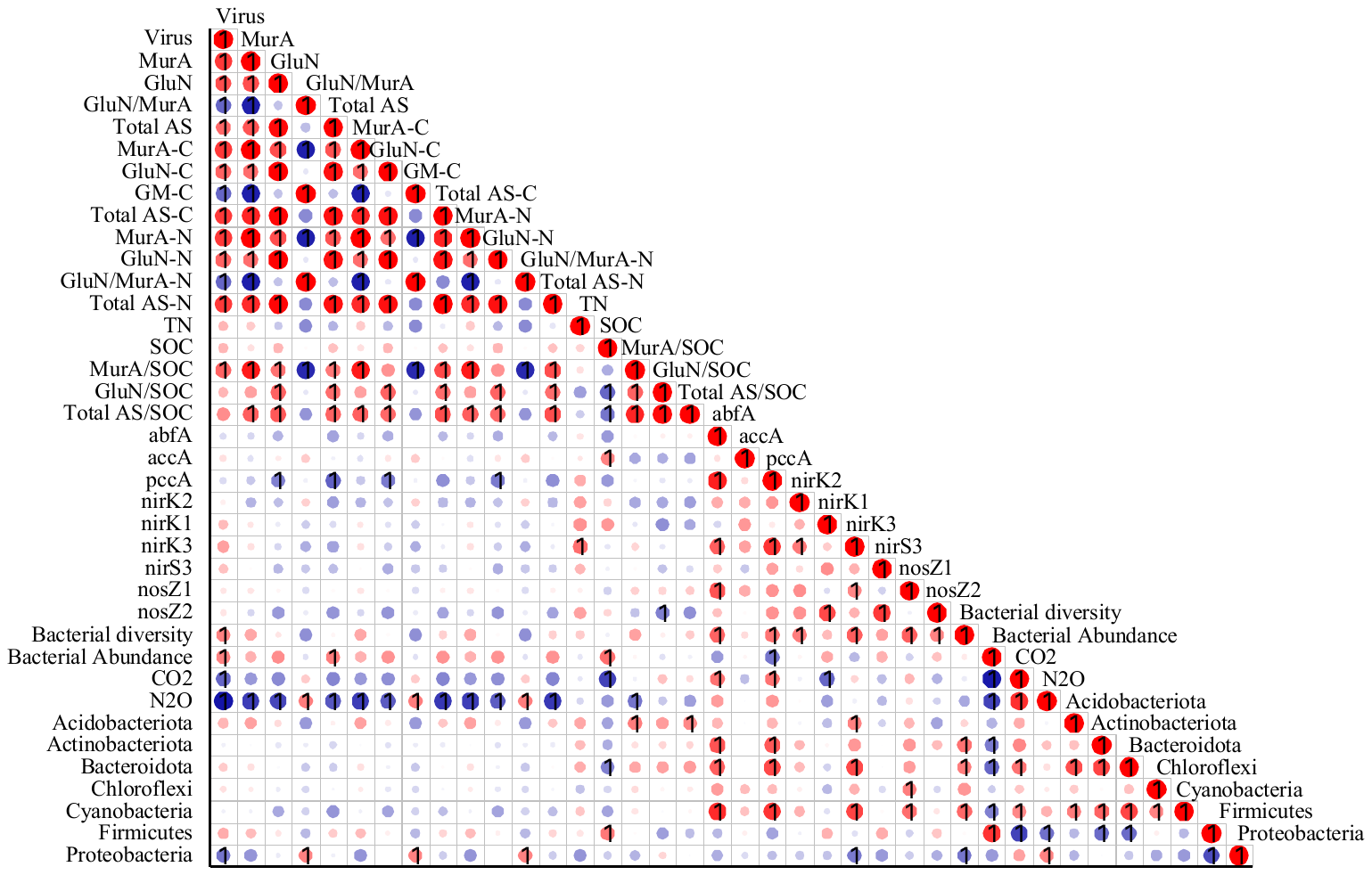

**Fig. S5** Pearson correlation analysis between viral addition, microbial community traits, soil physiochemical properties, greenhouse gas emissions and SOC sequestration. Red color indicates positive correlations, and blue color indicates negative correlations. The number 1 suggests correlation of statistical significance.

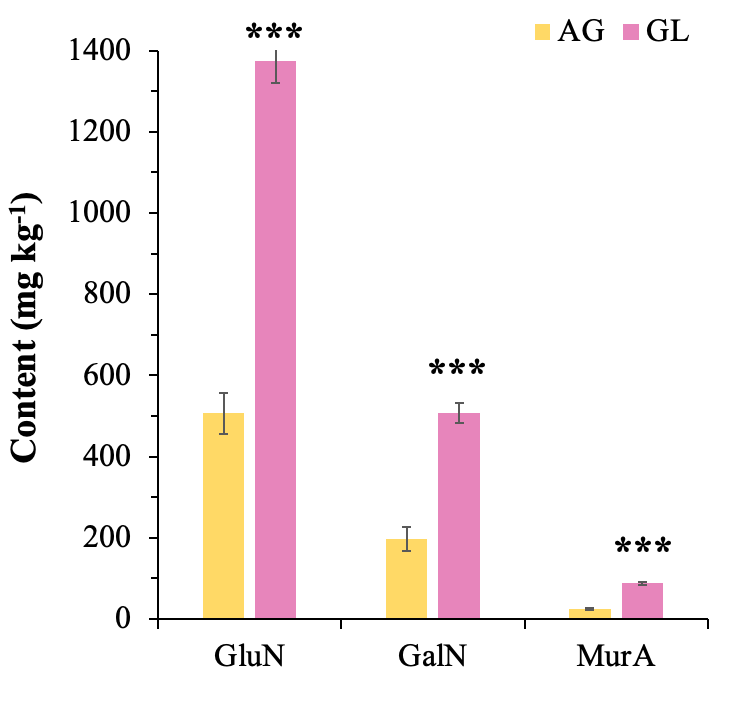

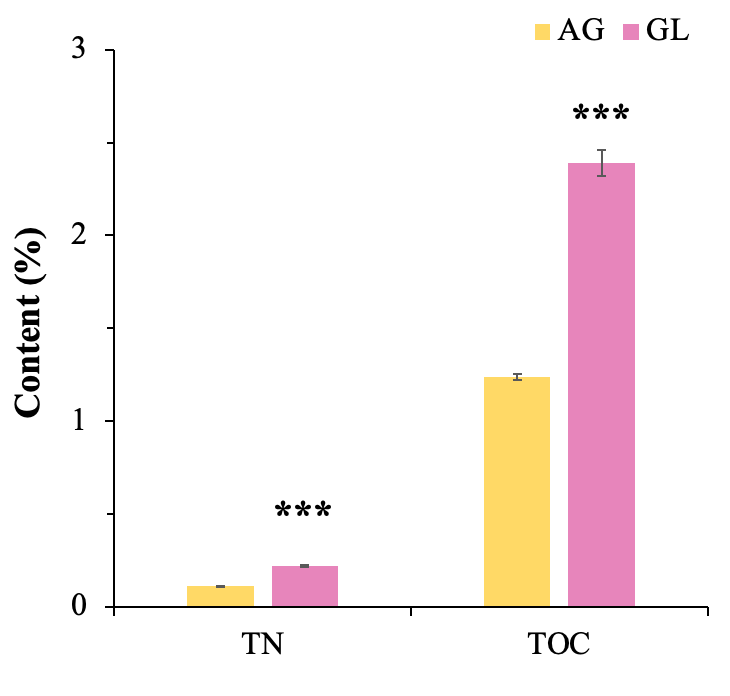

**Fig. S6** The content of amino sugars, i.e., muramic acid (MurA), glucosamine (GluN), and galactosamine (GalN), total organic C (TOC), and total nitrogen (TN) in the agricultural (AG) and grassland (GL) soils.
